# The Frequency of Population and Masting Fluctuations Is Increasing Globally

**DOI:** 10.64898/2026.02.04.701112

**Authors:** Vincent S. Pan, Peter E. Rothstein, Kadeem J. Gilbert

## Abstract

Substantial anthropogenic changes to the environment have motivated efforts to quantify temporal trends in population dynamics. While most ecological research has focused on the mean and variance of population density and reproduction, the frequency of these fluctuations through time may also be changing. We analyzed 1,563 datasets of population density and 1,456 datasets of plant reproduction (masting) across the globe. The average frequency of fluctuations increased by ∼ 0.5 – 3% per decade within each time series, representing a moderate change (Cohen’s *d* ≈ 0.4) over a period of 60 years. We tested four hypothesized mediators of this trend: increased temperature, increased frequency of environmental forcing, increased intrinsic growth rate, and increased distance from a saddle at zero density. Although all hypotheses were rejected, changes in the frequency of environmental forcing and intrinsic growth rate exhibited positive correlations with changes in population fluctuation frequency as expected. Our results suggest that successive peaks in population and masting density fluctuations are becoming closer in time, which may reduce the effectiveness of predator satiation, resilience of food-webs, and the risk of critical transitions, such as population extinction. We suggest some alternative hypotheses for what may underlie this surprising global pattern.

## Introduction

Anthropogenic changes to the environment have led to conspicuous changes to biodiversity (Wagner 2020). Understanding how population dynamics respond to global change is therefore a major goal of 21^st^ century ecology. But whereas much research has focused on understanding changes in the mean (Edwards et al. 2025, Johnston et al. 2025) and variation (Pearse et al. 2017, Gilarranz et al. 2022) of population density, dynamics can vary in multiple aspects. Characterizing changes in different aspects of dynamics is a first step in distinguishing between competing hypotheses of what governs dynamics (Kendall et al. 1999, Dwyer et al. 2004) and how they might change in the future. Accordingly, we tested whether an often-overlooked aspect of population dynamics—the inter-annual frequency at which population density fluctuates—has been changing globally across diverse taxa. We also examined reproductive dynamics as they can generate population cycles (Myers 2018) and may be especially sensitive to climate change (Hacket-Pain and Bogdziewicz 2021). A change in frequency, if it exists, has important implications for population persistence (Inchausti and Halley 2003), regime shifts (van der Bolt et al. 2018), and the resilience of food-webs (Yang et al. 2019).

Frequency may be changing for several reasons: (i) Declines in population growth rates documented across diverse taxa (Buckley et al. 2010, Edwards et al. 2025), if widespread, correspond to a decrease in population fluctuation frequency (Ricker 1954, Rosenzweig and MacArthur 1963). (ii) Likewise, declining global climate annual fluctuation frequency (Boulton and Lenton 2015, Di Cecco and Gouhier 2018) implies that populations (Pepi et al. 2021) and reproduction (Ascoli et al. 2021) influenced by climate forcing may show similar declines in fluctuation frequency. (iii) Conversely, rising global temperatures (Hansen et al. 2010) may accelerate biochemical reactions and metabolic theory predicts an approximately exponential decrease in physiological time (Gillooly et al. 2001) and population cycling period (Peterson et al. 1984). Similar hypotheses have been developed for plant reproduction under the assumption that warmer temperatures accelerate photosynthesis (Bogdziewicz et al. 2020), making conditions suitable for reproduction more frequent (i.e., fewer ‘vetos’ *sensu* Pearse et al. 2016) or resource accumulation faster between successive seeding events (Satake and Bjørnstad 2008). (iv) Finally, as a population’s density fluctuates, the density at which the population trajectory approaches a saddle at zero density may change. For example, a population may experience stronger predator suppression at densities close to zero as landscape simplification promotes predator foraging efficiency or preference (Kamaru et al. 2024). Theory indicates that trajectories sufficiently close to a saddle will remain transiently in the saddle’s vicinity (‘saddle crawlby’ *sensu* Hastings et al. 2018, Rubin et al. 2023), making population cycling frequency proportional to distance from the saddle. These mechanisms suggest that fluctuation frequencies in population and reproductive dynamics may be shifting in the Anthropocene yet remain underexplored.

Here, we investigate how and why the frequency of population and reproduction fluctuation has been changing globally over roughly the past two centuries. We analyzed population (1,456 datasets, 474 species) and reproduction (1,563 datasets, 477 species) time series in the Global Population Dynamics Database (Figure 1 A-C, Prendergast et al. 2010) and MASTREE+ database (Figure 1 D-F, Hacket-Pain et al. 2022) respectively. We focused on masting, the highly variable and spatially synchronous reproduction of plants, because its dynamics are strongly shaped by weather conditions (Pearse et al. 2016) and its creation of pulsed resource fluxes strongly shape ecosystem dynamics (Michaud et al. 2024). Finally, we tested whether mechanisms (i) – (iv) and (ii) – (iii) could be mediators explaining a shift in population and masting fluctuation frequency respectively.

**Figure 1.**
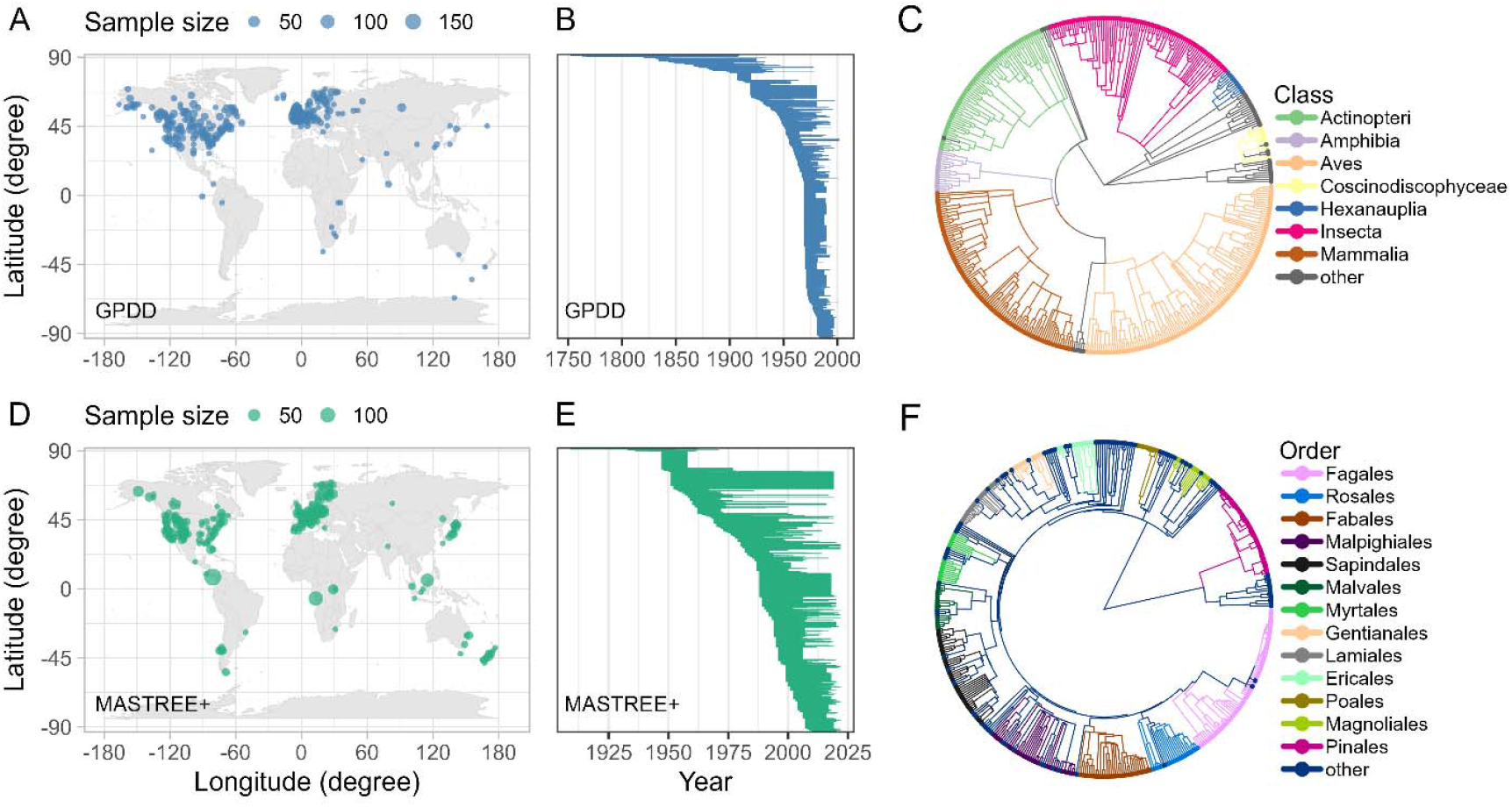
The sampling extent of (A-C) population and (D-F) masting density time series. We included only series of population or reproductive density of single species that are at least eleven years long with less than 20% of the data missing. Data for each series are standardized on an annual basis by finding the yearly maximum (though the results remain robust to using the yearly mean). In (A, B), GPDD stands for the Global Population Dynamics Database. (A, D) The geographic distribution of census locations binned at 2 □ × 2 □. Point size is scaled relative to the number of time series. (B, E) The temporal extent of time series analyzed. Each line segment denotes the time span of a single time series. (C, F) The phylogenetic sampling of the time series. Taxonomic classification is gathered using *taxadb* (Norman et al. 2020) and *taxizedb* (Chamberlain et al. 2023). In the legends, clades with fewer than ten sampled species are displayed as ‘other’. In (C), the phylogeny is pruned from the most recent Open Tree of Life (Hinchliff et al. 2015) synthesis tree (package *rotl* Michonneau et al. 2016), with branch lengths not to scale. In (F), the dated phylogeny is pruned from the most recent Angiosperm megatree (package *rtrees* Smith and Brown 2018, Li 2023).

## Methods

### Global trend of frequency

To estimate the temporal trend in frequency without bias from confounders that vary *between* series (e.g., life history differences), we compared the time-localized frequency spectra *within* each time series. We estimated time-localized frequency spectra by cutting each series into two otherwise identical segments from the center (Figure S1A) and detrended each log-transformed segment with LOESS before computing the Lomb frequency spectrum (Lomb 1976, Figure S1B, on average resolving frequencies between ∼ 0.1 – 0.5 cycles / year). For each spectrum, we computed the geometric mean frequency weighted by the linearly interpolated power of each frequency.

We estimated the average rate of change in mean frequency across all masting or population time series by fitting the mean frequency of each segment against the center year of each time series and the offset year of each segment from the series center (Figure S1C). The slope associated with the offset variable is the average marginal rate of change in log frequency *within* each time series, the quantity used to test our hypothesis. This method is equivalent to partitioning trait correlation *between* and *within* species used in phylogenetic comparative analysis (de Villemereuil and Nakagawa 2014) and the ‘Econometric Fixed Effects Design’ *sensu* Byrnes and Dee (2025). We used Bayesian generalized linear mixed models (GLMM) with moderately regularizing priors (package *brms*, Bürkner 2017). We used a log-linked gamma distribution as the conditional distribution and included the log series length as a fixed effect.

Census location, study identity, and series identity were included as random effects in all models. For the analysis of population density, we included serially nested species, family, and class to account for species relatedness as no phylogeny with branch lengths is available. For the analysis of masting density, we included masting variable unit nested within variable type, species, and the phylogeny as random effects.

### Mediators

We investigated whether mechanisms (i) – (iv) could have mediated the temporal trend in frequency. For intrinsic growth rate, we fitted the stochastic Ricker model to each segment (Ricker 1954, Delean et al. 2013, see supplement). For temperature, we obtained reconstructed yearly mean temperature at each census location from 1806 – 2024 (Fan and van den Dool 2008, Slivinski et al. 2019) and computed the mean inverse temperature for each segment. The inverse transformation follows from the expectation that the log frequency is roughly proportional to the inverse absolute temperature in the Arrhenius equation (Gillooly et al. 2001). For environmental forcing frequency, we used temperature as a stand in for the environmental driver and computed the mean temperature frequency using the same method described above. For distance from saddle at zero, we used the log minimum observed density of each segment.

Mediation was estimated by fitting multivariate Bayesian GLMMs with the same fixed and random effects as above, but with the center and offset of mediators also included as fixed effects. The offset of each mediator was also fitted against the offset of segment year in their respective sub-models along with the same random effects as the frequency sub-model. We used an identity-linked normal distribution as the conditional distribution for each mediator. Mediated effects were computed by multiplying the posterior distribution of the path coefficients from segment year offset to frequency. Analyses were performed in R v4.4.2 (R Core Team 2025).

## Results

We found that frequency of population and masting density fluctuations has been increasing by 1.32% (95% credible intervals = [0.818, 1.83]%, ΔR^2^ = 0.018 [0.0065, 0.034]) and 1.1% ([0.585, 1.62]%, ΔR^2^ = 0.014 [0.0036, 0.029]) per decade on average within each time series respectively (Figure 2A, C). These results are mostly robust to different data inclusion criteria and are not a result of statistical artifacts, with longer and more cyclic time series showing roughly twice the rate of increase than the typical time series (Figure 2B, D). The rate of increase for cyclic masting series yielded a similar estimate but is indistinguishable from zero, likely due to the small sample size. Typical population cycle period shows a standard deviation of ∼ 30% of the mean (Dwyer et al. 2004, Rubin et al. 2023), placing our estimated trend at ∼ 0.017 – 0.10 Cohen’s *d* per decade across models (Figure 2B, D) and indicating that the magnitude of change is small ecologically over the short-term. Although whether the trend may continue over longer terms remains unclear, our analysis of only long time series (Figure 2B, D) revealed that the frequency of population (*d* = 0.44 [0.22, 0.65]) and masting (*d* = 0.40 [0.25, 0.55]) density fluctuations has already increased moderately within 60 years, the average length of the time series.

**Figure 2.**
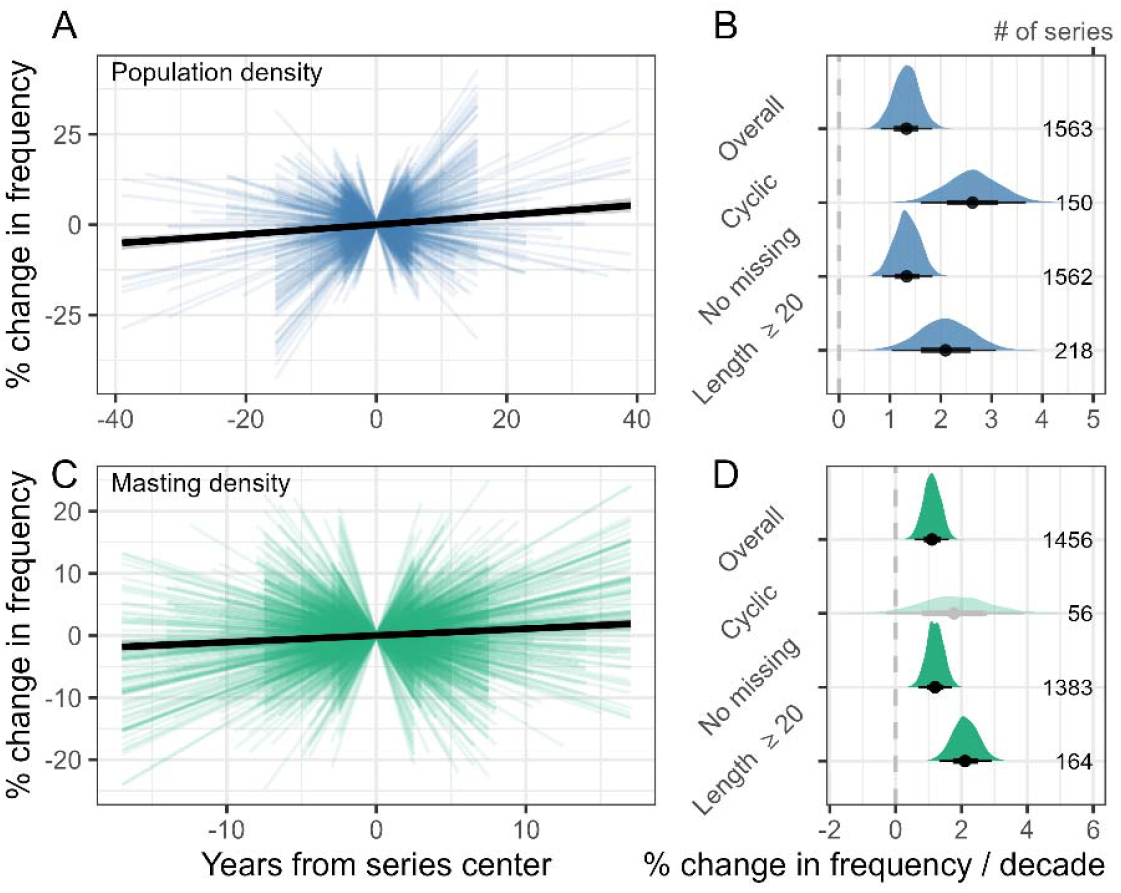
The frequency at which (A-B) population density and (C-D) masting density fluctuate is increasing through time within each time series. (A, C) The relative change in frequency as a function of time, with the center point of each time series as the reference. Thick black lines and ribbons show the posterior mean and 95% credible intervals of models fitted to the full respective datasets. Thin colored lines show the raw relative change for each time series. (B, D) As robustness checks, we refitted the models to data subset by three criteria. (1) We examined only cyclic time series because the fluctuation frequencies in these series often inform mechanisms of population fluctuation (Murdoch et al. 2002, Dwyer et al. 2004). We classified whether a series is cyclic conservatively via the false-alarm-probability of the dominant frequency at a type I error rate of 5% (Press 2007). (2) We examined only series with no missing data because differences in the degree of missingness between segments may bias the estimation of the change in frequency. (3) We examined only long series (20+ years per segment) because the frequency resolution and frequency recovery accuracy are much higher. Posterior distributions of the rate of change according to these criteria are displayed. Points, thick lines, and thin lines show the mean, 66%, and 95% credible intervals. Distributions with 95% credible intervals that overlap zero have a higher transparency. The number of series analyzed is displayed as a number on the right.

We found that none of the hypothesized mediators could explain the observed increase in frequency (Figure 3A-H). Changes in population fluctuation frequency increased with changes in population growth rate and temperature fluctuation frequency as expected (Figure 3A, C), but both growth rate (-0.038 [-0.084, 0.0064] year^-1^ / decade) and temperature fluctuation frequency (-23 [-27, -20]% / decade) may have decreased with time, thereby mediating a decrease, not an increase, in frequency (Figure 3D). Conversely, temperature (0.22 [0.18, 0.27] K^-1^ / decade) and distance from saddle (21 [16, 27]% / decade) have decreased and increased with time respectively but neither showed the expected relationship with changes in population fluctuation frequency (Figure 3 B, E) and nor mediated an increase in frequency (Figure 3D). Finally, for masting plants, we found that although the temperature (-1.1 [-1.1, -1.0] K^-1^ / decade), not the fluctuation frequency of temperature (-1.9 [-7.1, 3.5] % / decade), has increased on average, increasing temperature was not associated with an increased frequency of masting (Figure 3F). Thus, both variables mediated an effect that is indistinguishable from zero (Figure 3H).

**Figure 3.**
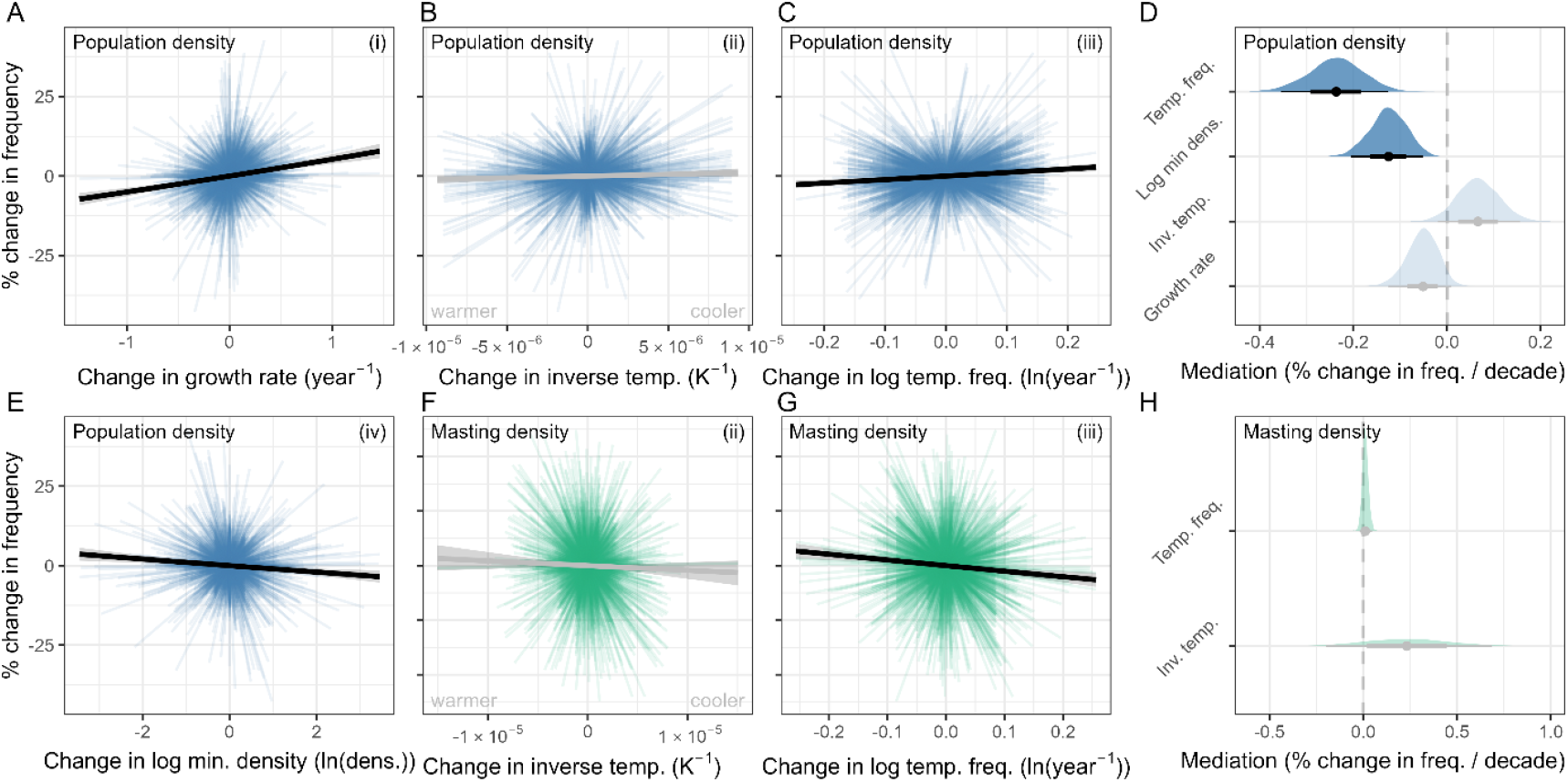
None of the hypothesized mediators tested could explain the observed increase in frequency of fluctuation in (A-E) population density and (F-H) masting density. (A-C, E-G) The relative change in frequency as a function of each mediator, with the center point of each time series as the reference. Thick black lines and ribbons show the posterior mean and 95% credible intervals. The lines are greyed out if the 95% credible intervals overlap zero. Thin colored lines show the raw relative change for each time series (population density: 1,535 series; masting density: 1,456 series). The numbered hypotheses described in the text are labeled on the top right corner. (D, H) Posterior distributions of the indirect effects of time on frequency mediated through different variables. All indirect effects were hypothesized to be positive, falling to the right of the dashed line. Points, thick lines, and thin lines show the mean, 66%, and 95% credible intervals. Distributions with 95% credible intervals that overlap zero have a higher transparency.

## Discussion

We have shown that the fluctuations in population and masting density are becoming more dominated by high-frequency oscillations at an average rate of 0.5 – 3% per decade across diverse taxa globally. While negligible in the short-term, ecologists monitoring a population over a lifetime would have observed on average a modest increase in frequency, as is indeed the case (Övergaard et al. 2007, Pesendorfer et al. 2020, Shibata et al. 2020). This result has profound implications for population and community dynamics, as well as our management of these systems. While decreased climate fluctuation frequency has led to concerns that populations may be more vulnerable to extinction (Di Cecco and Gouhier 2018) and irreversible regime shifts (van der Bolt et al. 2018), our result suggests a more optimistic outlook as population fluctuation frequency increased rather than decreased. However, the dynamics of these populations may be harder to predict and their effect on their surrounding community may be less optimistic. Indeed, for species that are dependent, but not driving the dynamics of another fluctuating species, theory suggests that they may be less resilient to perturbations (Yang et al. 2019) and less likely to coexist (Miller and Klausmeier 2017). Seeds that depend on predator satiation for survival may face increased mortality as predator populations remain high between mast years, a possibility supported by a recent meta-analysis (Zwolak et al. 2022).

Surprisingly, none of the hypothesized mediators explained the increase in frequency; rather, it increased despite these mediators, suggesting that other unmeasured drivers of fluctuations exist and may be changing. The striking similarity between the rate of increase in masting and population fluctuation frequency, while perhaps coincidental, intriguingly suggests a common driver. What might these drivers be?

We suggest three alternative hypotheses. First, the change in frequency may be driven by an increase in forcing frequency of other environmental factors besides temperature, including precipitation (Pepi et al. 2021) and the difference between inter-annual temperatures (Δ*T sensu* Kelly et al. 2013). Second, global eutrophication (Gruber and Galloway 2008) can release populations from resource limitations, shifting low-frequency consumer-resource oscillations to high-frequency density-dependent oscillations (Murdoch et al. 2002). For masting trees released from nutrient limitation, resource-budget models predict shorter inter-masting periods (Satake and Bjørnstad 2008). Finally, the increase in frequency may be an intrinsic and universal nature of population and masting dynamics irrespective of global change. Tree fecundity can change with age (Qiu et al. 2021) leading to changes to masting frequency (Pesendorfer et al. 2020). Populations can also ‘age’ as processes slower than population dynamics evolve and shorten cycle period. These processes may include local adaptation and spatial dynamics (e.g., self-thinning Westoby 1984) that take much longer to reach equilibrium. Because the increase in frequency appears to be widespread and important, we believe that testing these hypotheses is a fruitful way forward.

## Conflicts of Interest

We declare no conflicts of interest.

### Acknowledgements

We thank Sylvie Martin-Eberhardt, Nasser Rabi, Ian Pearse, Daniel Mok, Cristal Lopez, and Mataeus Funderburk for helpful discussions. This is Kellogg Biological Station contribution no.???

## Data archiving statement

All code and data are deposited at Zenodo (https://doi.org/10.5281/zenodo.18301800) and GitHub (https://github.com/vsbpan/specter). A zipped repository file is attached as a supplement for review. The DOI will be live upon manuscript acceptance.

## Author contributions

Conceptualization: VSP, PR; Formal analysis: VSP, PR; Methodology: VSP, PR; Software: VSP, PR; Writing – original draft: VSP; Writing – review & editing: PR, KJG; Supervision: VSP, KJG

## Supplemental Materials

### Ricker model estimation

Writing *Y*_*t*_ as the log density at time *t* ∈ {2,…,*T*}, *r* as the intrinsic growth rate, *K* as the carrying capacity, and *ϵ*^2^ as the residual variance, we estimated,

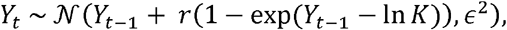

using maximum *a posterori* with informative priors for each population time series segment. For priors, we set,

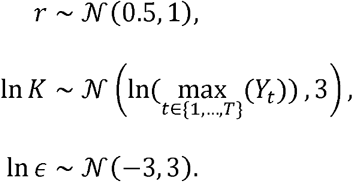

### Computation of ΔR^2^

We computed ΔR^2^ as a measure of the proportion of variance of the frequency *within* series that is explained by the change in time. Denoting the frequency of a segment *j* ∈ {1,…, *J*} of time series *i* ∈ {1, …,*n*} as *f*_*i,j*_, the observed proportional change *within* series is 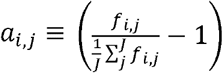. Using the estimated slope 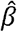 of the center year offset variable *x*_*i,j*_, we define,

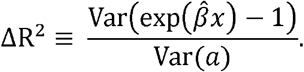

**Figure S1.**
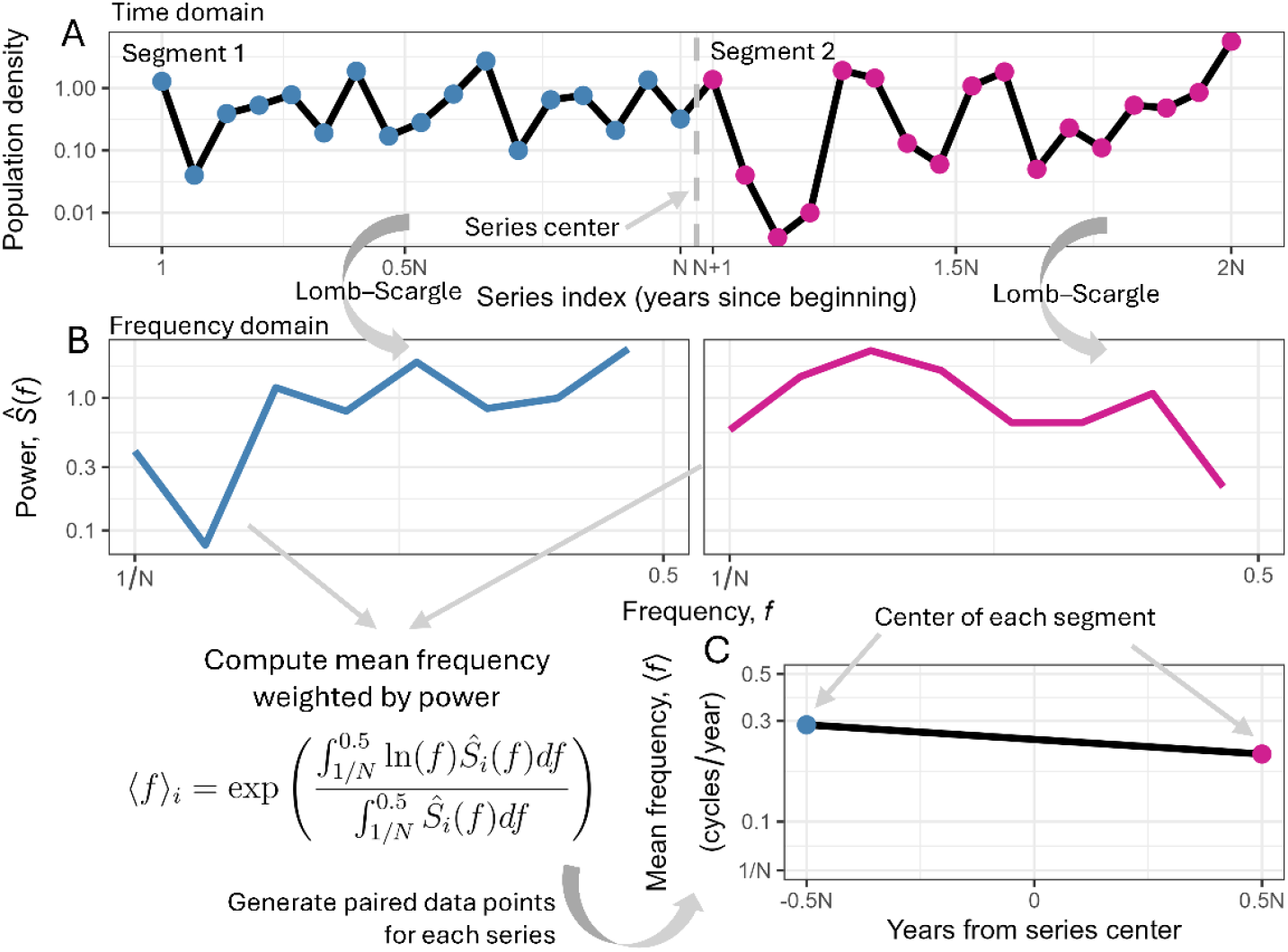
A schematic diagram of the approach we took to estimate the average rate of change in frequency within each time series. (A) A raw time series of length 2*N* in the time domain is cut into two segments of length *N* from the center. The data show the population density through time of *Arctia virginalis* caterpillars at Bodega Bay Marine Reserve from 1986 to 2019 (Pepi et al. 2021). (B) Each time series segment is log-transformed, detrended, and transformed to the frequency domain via the Lomb–Scargle method, which can deal with missing data. For a segment of time series of length N, we can resolve ⌊ *N*/2 ⌋ frequencies from 0.5 to 1/*N* cycles per year. The obtained spectra show good agreement with the analysis of Pepi et al. (2021) using wavelet analysis to time-localize frequencies (dominant frequency ∼ 0.3 – 0.5 cycles / year between 1992 – 1999; 0.17 – 0.25 cycles / year between 2003 – 2013). (C) The continuous frequency spectrum 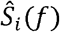 of each segment *i* is obtained by linear interpolation between the power spectral density of each resolved frequency and used to compute the weighted mean frequency. We assumed that the mean frequencies are time-localized at the center of each segment. The series center year and segment year offset (segment year - series center) are used as predictors to partition time correlation *between* and *within* time series. The pair data points show qualitative agreement with the findings of Pepi et al. (2021) that altered precipitation dynamics in Northern California led to a regime shift in *A. virginalis* dynamics starting around 2004 (*N*+2).

